# Paired-end Mappability of Transposable Elements in the Human Genome

**DOI:** 10.1101/663435

**Authors:** Corinne E Sexton, Mira V Han

## Abstract

Though transposable elements make up around half of the human genome, the repetitive nature of their sequences makes it difficult to accurately align conventional sequencing reads. However, in light of new advances in sequencing technology, such as increased read length and paired-end libraries, these repetitive regions are now becoming easier to align to. This study investigates the mappability of transposable elements with 50bp, 76bp and 100bp paired-end read libraries. With respect to those read lengths and allowing for 3 mismatches during alignment, over 68%, 85%, and 88% of all transposable elements in the RepeatMasker database are uniquely mappable, suggesting that accurate locus-specific mapping of older transposable elements is well within reach.

## Background

Sequences from transposable elements (TE) have been observed to account for an average of 7% of reads from human RNA-Seq datasets (1). Because of the repetitive nature of TE sequences, fragments originating from locus-specific TEs are challenging to accurately align to a reference genome. However, RNA-Seq protocols now allow for paired-end reads up to 150bp long which theoretically should substantially increase the ability to confidently map to repetitive regions (2). Mappability of the genome has been analyzed before, and several representations have been proposed, including pair-wise cross mappability, and minimum unique length (3–6), but, no study has investigated the mappability of TE regions in light of differences in read lengths and paired-end sequencing. The purpose of our study is to assess which TEs can be confidently quantified at the locus-specific level, rather than at the aggregated family level. We provide base level mappability of TEs in detail and show that a significant fraction of TEs are uniquely mappable at the element level.

Mappability has been defined for the genome by Derrien et al as the inverse of the number of times a k-mer originating from the genome appears in the genome, considering its reverse complement and allowing a limited number of mismatches (5). We follow this definition to attain mappability scores for all TE regions in the RepeatMasker database using both the GEM (GEnome Multitool) Mappability and Bowtie programs (5,7).

Several factors affect the ability to accurately map a sequencing read to its originating locus which are not accounted for in this analysis. These include sequencing errors in the reads, true variants between the genome which was sequenced and the reference genome, and the accuracy of the alignment software used to map the reads. Both random sequencing errors and true genomic variants can potentially create sequencing reads that are not present in the genome (8,9). Such reads are not generated in the simulation in this analysis, but by allowing for up to 3 mismatches for each read in our alignment, we are capturing possible mappings to potential errors and variants up to 3 nucleotides for every sequence of read length in the genome. Also, since no alignment software maps sequencing reads perfectly (1), we compare our estimates of mappability based on two independent alignment algorithms, GEM and Bowtie, and check whether the estimates are largely congruent between the two methods.

Additional factors which affect the ability to discern the originating position of reads when looking at TE loci specifically include the copy number of the TE subfamily (10), the Hamming distance between related loci relative to the allowed mismatch rate and, importantly, the accuracy of TE annotations. In this analysis we use the Repeatmasker TE annotation database for all TE classifications, but the current lack of benchmarking methods for TE annotation likely results in missing annotations or incorrect annotations even in this database (11). Our analysis takes this fact into consideration as we have calculated mappability scores for the entire genome and therefore TE locus level scores would be simple to recalculate based on any future updates to TE annotations.

In our analysis, we provide a specific mappability score which represents the ability of Bowtie software to map a read pair uniquely to a locus when allowing for up to 3 mismatches. Although this is a very specific definition of mappability and we cannot account for all of the above listed confounding factors, these scores provide an estimate of the sequence uniqueness under particular parameters and assumptions (12). While mappability scores are not directly related to how well a read maps to a region, they do provide a conservative estimate of the uniqueness of a sequence which can inform on the resulting confidence of a read mapping.

We have provided six new hg38 mappability tracks for the UCSC browser as well as provided aggregate mappability scores for each TE locus found in the RepeatMasker database. We show that more than 85% of TE loci are considered to be unique in terms of 76bp paired-end mappability allowing for 3 mismatches using the Bowtie aligner. Because these TE loci have unique mappability scores, we present these TE loci as a high confidence set that researchers can utilize as a reference for studying TE expression based on RNA-seq.

## Methods

We used two approaches to estimate the mappability of repeat elements. First to obtain baseline single-end read mappability scores, we used GEM Mappability, an efficient software used to rapidly determine mappability with multiple mismatches. And second, we generated all possible k-mers from the reference genome sequence, and mapped them back to the reference genome using Bowtie reporting all mappings. We considered this approach would be complementary to GEM Mappability because the scores from Bowtie 1) represent how an aligner would actually map the reads and 2) can accurately assess paired-end read mappability by allowing for a range of insert sizes.

Additionally, Karimzadeh et al have more recently described Umap (4) as a method for scoring genome mappability. Though Umap does map kmers to the genome using Bowtie, similar to our method, we omit a comparison to this software because the definition of mappability used in Umap is vastly different from both GEM Mappability and our method. Umap defines mappability as “the fraction of a region that overlaps at least one uniquely mappable k-mer” whereas we define mappability as the inverse of the number of times a k-mer originating from the genome appears in the genome. In other words, our method depends on the start position of a kmer to determine mappability scores, whereas Umap takes into account the length of the kmer in its scoring.

Additionally, the default Umap mappability tracks were created using Bowtie not allowing any mismatches whereas our Bowtie mappability scores allowed for 3 mismatches. This results in dramatically different scores as well and for these reasons we believe a direct comparison between GEM Mappability, Umap and our method could be misleading.

### GEM Mappability Estimates

To get single-end mappability scores for comparison, GEM Mappability (build 1.315) (5) was run on hg38 with a read length of 50, 76, and 100 base pairs and an allowance of 3 mismatches.

### Read Generation for Bowtie Mapping

To represent all possible read sequences that could be transcribed from the reference genome, we generated 76-mers from hg38 using jellyfish (13). We assume that no splicing events occur in the transcription of the transposon sequence. Although, this assumption can be wrong for certain transposons, it is generally applicable to the majority of transposons. To simulate paired-end reads, we generated all possible 200-mers, 242-mers, and 300-mers from hg38 using jellyfish. For each of these sizes we removed a region between the two paired-end “reads,” (100bp, 90bp, and 100bp respectively) and reverse complemented the second “read” in each pair to simulate a forward-reverse paired-end library. In order to create a library similar to the RNA-seq data from the Genotype-Tissue Expression (GTEx) project (5) insert length of 242bp was chosen for the 76bp paired-end library based on the median insert length across GTEx bamfiles.

### Bowtie Mappability Estimates

Using Bowtie (v1.2.2) (7) with the -a all alignments flag, both the single 76-mer set and the three paired-ended 50-mer, 76-mer, and 100-mer sets generated above were mapped back to their originating reference allowing for 3 mismatches. We also investigated the effect of alignment by using STAR instead of Bowtie using options -- outFilterMultimapNmax 1000 --winAnchorMultimapNmax 100. Based on the preliminary analysis which showed Bowtie to generate generally more conservative mappability scores (Supplemental Figure 1), we decided to use Bowtie for the downstream analyses.

Throughout this article, when we refer to mappability of a single base position in the genome, it is based on the read alignments that start at that base position. We inferred the mappability of a position in the genome using the Bowtie alignments as follows. As an example, for a single base position, if a kmer mapped to that position and jellyfish reported that kmer to appear 5 times in the genome, the mappability of that position would be 1/5. Using the reported counts from jellyfish we can significantly simplify the process of mapping by including each unique kmer sequence only once in our read set while still retaining all pertinent information for the mappability calculation. Similarly, if multiple kmers map to a base position, as is often the case when considering mismatches, we calculate mappability by summing all reported jellyfish counts for each mapped kmer. For example, if 2 kmers map to a single position and jellyfish reports that one of the kmers appears 3 times in the genome and the other kmer appears only 1 time in the genome, the mappability for that base position would be 1/4.

An alternative, but equivalent, way of calculating mappability from a bamfile is to identify a kmer which maps exactly to a base position and take the inverse of how many times that kmer appears at other positions in the bamfile. For example, if a kmer maps perfectly to a base position and appears at exactly 4 other positions in the genome, then the mappability for that position would be 1/5. With this method, if 2 kmers mapped to the same base position, we would only consider mappability as the number of times the perfectly matching kmer appears in the bamfile.

These methods are theoretically equivalent, provided, jellyfish reports correct counts, and Bowtie reports the complete set of mappable positions. In reality, with single-end reads, we found that they are exactly equivalent (Supplemental Figure 2a). For paired-end reads, the two methods have a correlation coefficient of 0.98 with over 90% of mappability scores the exact same between the two methods (Supplemental Figure 2b). But the fact that the correlation is not 1 shows that there are some minor amount of errors in the approach that results in both missed mappings (points in the upper diagonal), or spurious mappings (points in the lower diagonal) in the Bowtie alignment, or vice versa for Jellyfish. The fact that we see more data points in the upper diagonal shows us that the method relying on Jellyfish counts (method 1) is more sensitive in retrieving all possible mappings compared to the method relying on Bowtie only (method 2). We use method 1 in our mappability calculations throughout this study.

### Transposon Locus Mappability

Extending this definition, we describe transposon locus mappability. If all positions in a transposon locus are uniquely mappable then the locus is considered uniquely mappable. Otherwise, if at least one position is not uniquely mappable, then the entire locus is classified as not uniquely mappable. We introduce this stringency because we are interested in a conservative mappability estimate. This approach also avoids possible mappability score inflation caused by unique flanking regions around TEs because the entire sequence must be considered unique for a TE locus to be classified as uniquely mappable.

## Results and Discussion

### Mappability Estimates across Software and Paired-end Results

In general GEM Mappability and Bowtie mapping scores have highly concordant results for single end mapping (Supplemental Figure 3), suggesting that Bowtie does indeed recover most if not all possible mapping locations when using the -a parameter. Since GEM Mappability software is also based on an independent alignment algorithm, the congruence between the two methods, especially for the exact matches, shows that the impact of alignment errors are limited. There are more discrepancies between the two methods as more mismatches are allowed, and it gives us some idea of the errors originating from mis-alignments. Even with 2 mismatches, the difference in mappability is less than 0.25 for 99.7% of the TE loci, and the correlation between the two methods is 0.98. We report the GEM Mappability results to represent single-end mappability going forward.

Figure 1 shows the mappability scores for both single-end and paired-end mapping. Each point in Figure 1 represents a TE with its mappability score generated from the 76bp reads with single-end (GEM) on the x-axis and paired-end (Bowtie) on the y-axis. The clear trend is an increase in mappability score with paired-end alignment. However, some paradoxical points have higher single-end mappability than paired-end mappability. These points make up less than 1% of total TE loci and can be accounted for by the presence of valid multiple mappings for the mate in the read pair (Figure 2).Especially in repetitive TE sequences, paired-end reads can map to the same starting position with varying insert sizes. This lowers the overall mappability score for a base position, in some cases making it lower than single-end mappability. An example of this is the AluY element at chrY:9192414-9192738. This 325bp-long element has 55 base positions where a varying insert size of the paired-end reads results in lower 50bp paired-end mappability scores compared to 50bp single-end mappability scores. It is relatively rare that varying insert sizes impact mappability scores in this way and therefore in analysis we treat differing insert size mappings conservatively as distinct mappings as shown in Figure 2.

**Figure 1:**
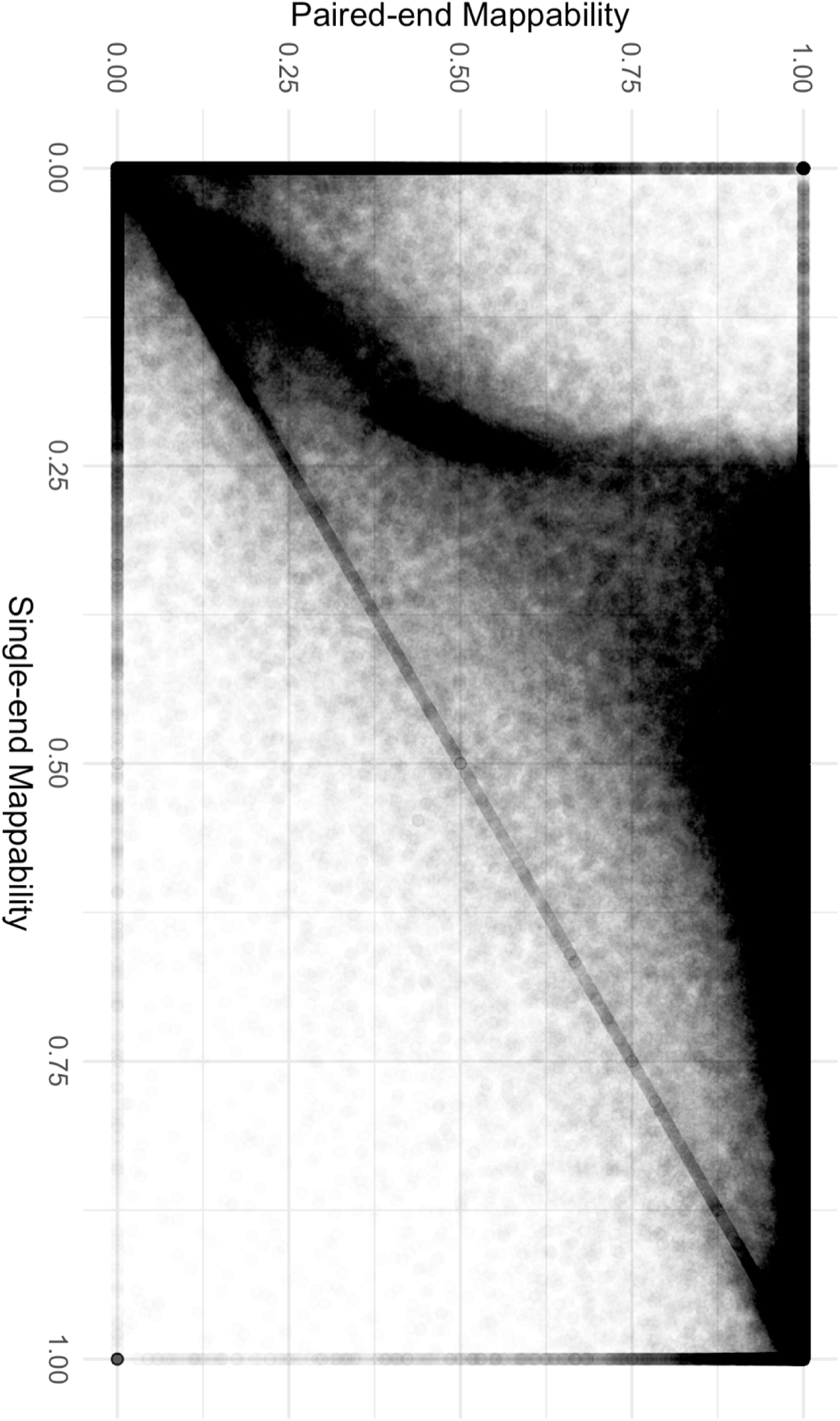
Comparison of single-end and paired-end TE mappability. Each point represents 1 TE sequence. Paired-end mappability scores were generated from 76 base pair read alignments with <= 3 mismatches. Single-end mappability scores were generated from GEM Mappability (k=76) with the default of a 4% mismatch rate. Mappability scores were aggregated for each TE as the percent of base pairs in the sequence with a unique mappability score.

**Figure 2:**
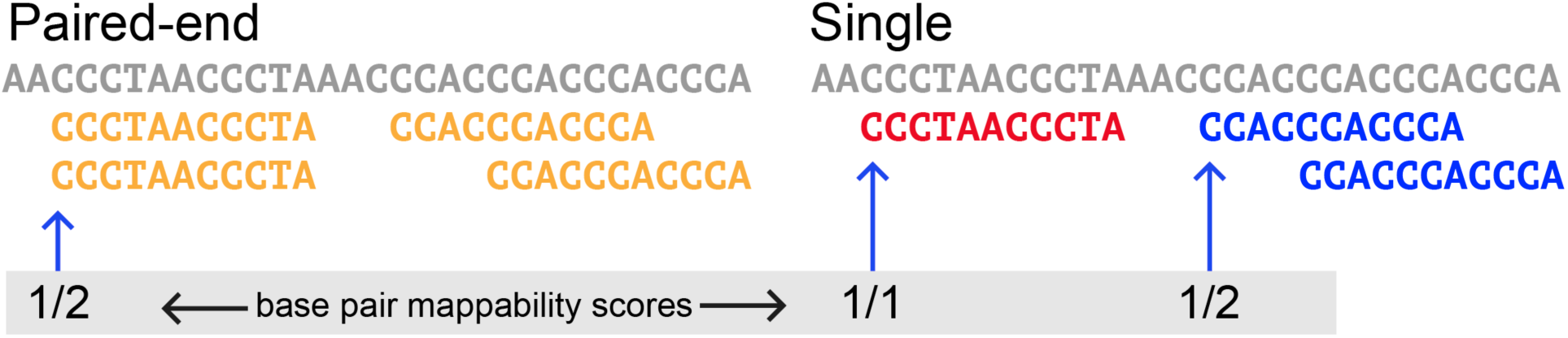
Paired-end and single-end alignment. Paired-end mappability for certain base pair positions may be lower than for the single-end mappability due to the read pair aligning multiple times in the same start position with a different insert size. In that case, the paired-end mappability score for that specific base position could be lower than the single-end mappability as in this example.

### Mappability of Elements

To assess general mappability for the TE sequences in the RepeatMasker database, we calculated both single and paired-end mappability scores as described in the methods. Figure 3 shows the percent of loci in each subfamily that are uniquely mappable for all three read library lengths in both single-end and paired-end configurations. We define a locus as uniquely mappable if all base positions at that locus had unique mappability scores.

Across all 4.67 million TEs defined in the Repeatmasker database, when allowing for 3 mismatches using the Bowtie aligner, 68.2% of all TEs are uniquely mappable with 50bp paired-end reads, 85.1% of all TEs are uniquely mappable with 76bp paired-end reads, and 88.9% of TEs are uniquely mappable with 100bp paired-end reads. At the class level using 76bp paired-end reads, 82.8% of SINE elements, 86.1% of LINE elements, 91.4% of DNA elements, and 84.7% of LTR elements are uniquely mappable (For full mappability statistics see Supplemental Table 1).

**Figure 3:**
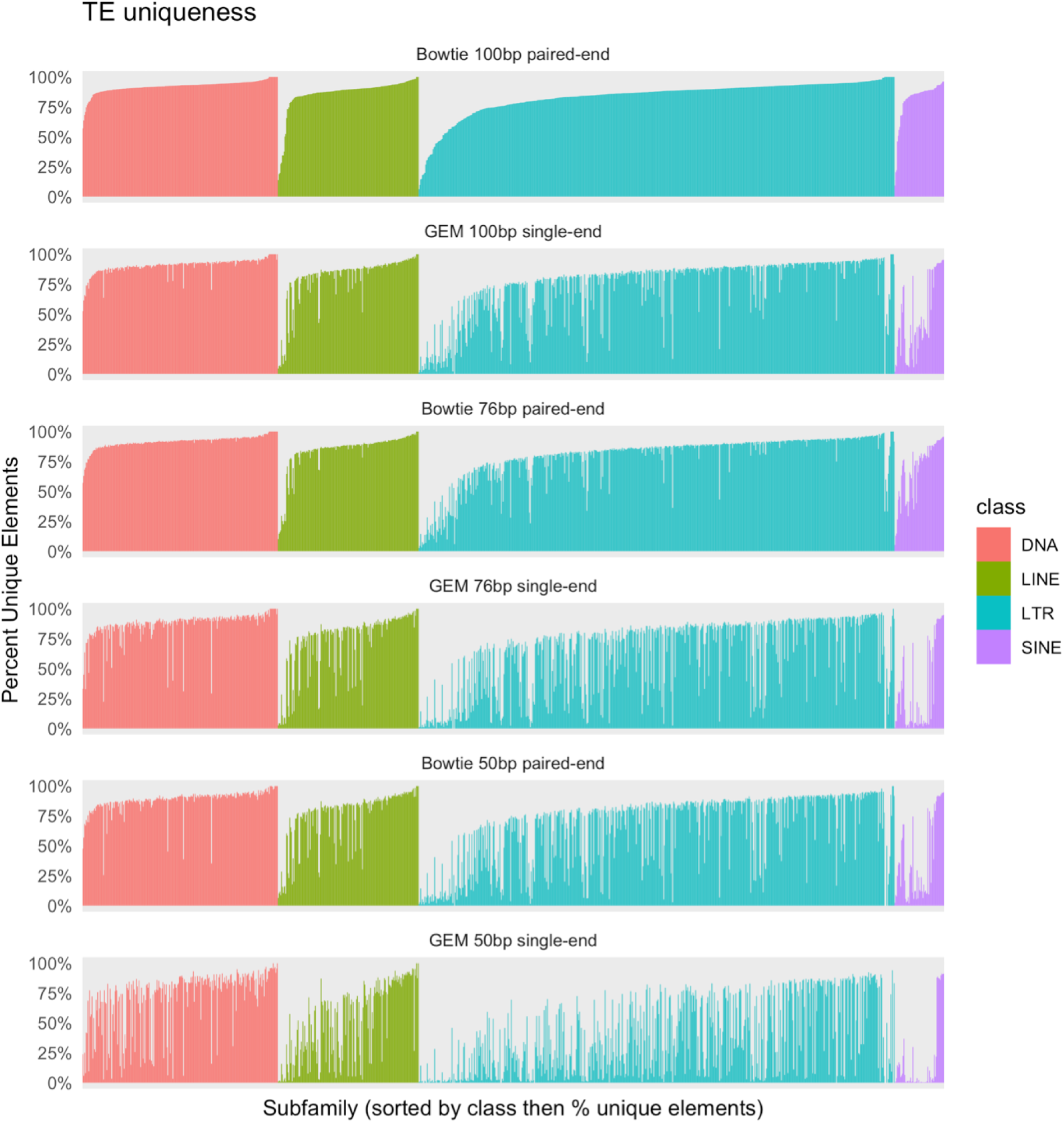
Percent mappable elements for each family and subfamily of TEs. Colored by class distinction, each bar represents the percent unique elements in each subfamily. An element is only considered unique if every position in its sequence has a unique mappability score. Both GEM Mappability and Bowtie were run with an allowance of 3 mismatches.

Given the nature of our mappability score calculation we would expect trends to arise with regards to copy number and age of TEs. More specifically, we expect as copy number for each subfamily increases, observed mappability scores should decrease due to multiple similar sequences being present in the genome. We investigated this relationship for each locus using 76bp paired-end mappability scores in Figure 4a. In general, the trend, though weak, is as predicted; as subfamily copy number increased, the average unique mappability scores for TE loci decreased. This trend may not be as strong as expected because the age of the TEs overshadows the effects of copy number; larger copy numbers do not necessarily lead to lower mappability, because mappability is also influenced by the mutations that have independently accumulated in each copy.

**Figure 4:**
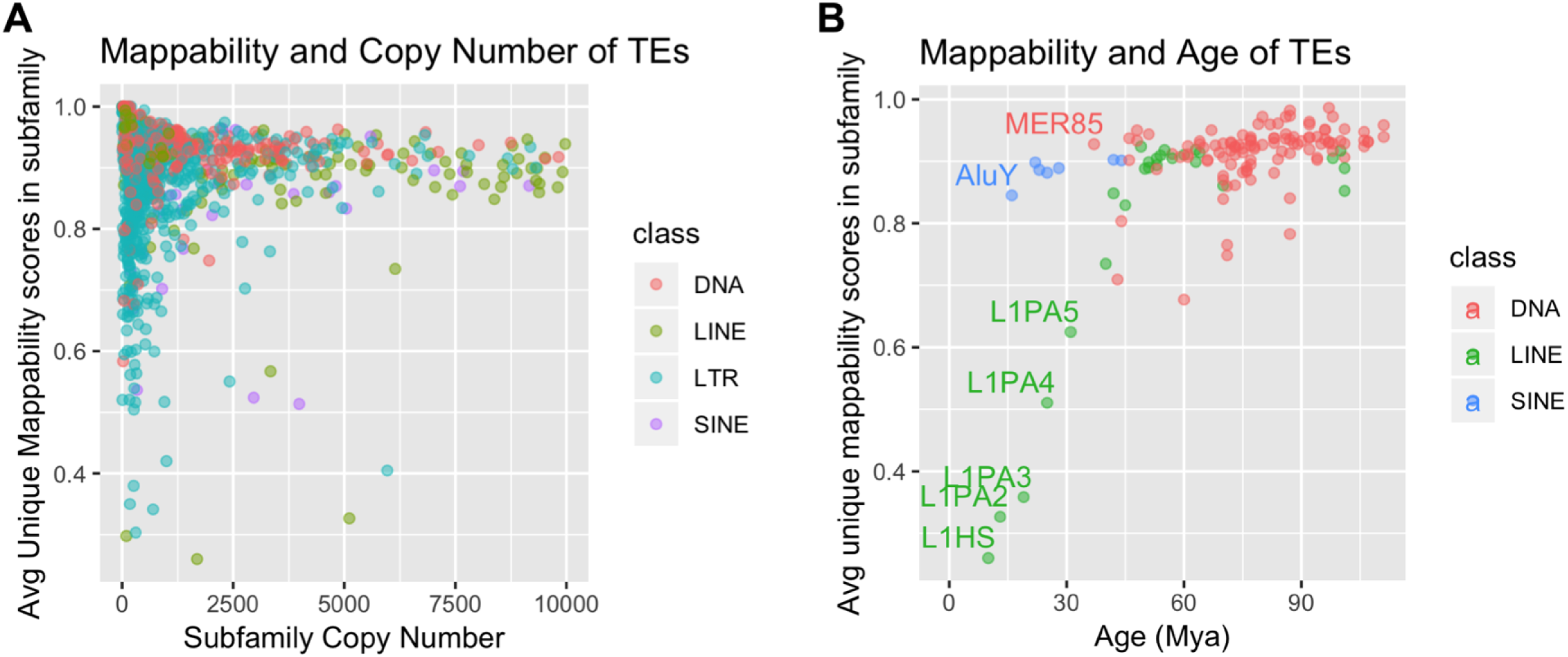
Mappability scores with regards to age and copy number of TEs. Plots of average unique mappability score in each TE subfamily colored by class. The average unique mappability score for each subfamily is calculated by averaging each individual locus’s percent unique sequence. The percent unique sequence is defined as the percent of uniquely mappable base positions in each individual locus. These aggregated scores are plotted against (a) copy number and (b) age for each subfamily. The ages in (b) were predicted in Pace and Feschotte (14).

To examine age and its effect on mappability scores, in Figure 4b, we plot the average unique 76bp paired-end mappability scores for TE subfamilies with ages predicted in Pace and Feschotte (14). This trend is much clearer; as age decreases, mappability score decreases as well.

Specifically, the families with the least mappable elements include some of the youngest, namely L1HS, L1PA2, L1PA3, AluY, and AluYa5. It is well known that the L1 and Alu families both contain elements which are considered to be active transposons (15,16). Because identical copies have recently been inserted in the genome, this decrease in mappability is expected. However even for these low-mappability elements a substantial increase in mappability is achieved on average with paired-end reads when compared to single-end 50bp reads (Figure 5). There are a few data points where the paired end mappability is lower than the single ended mappability, due to the issue of variable insert sizes described above. The outlier with the lowest value in AluY in Figure 5 is the AluY element at chrY:9192414-9192738 described above.

**Figure 5:**
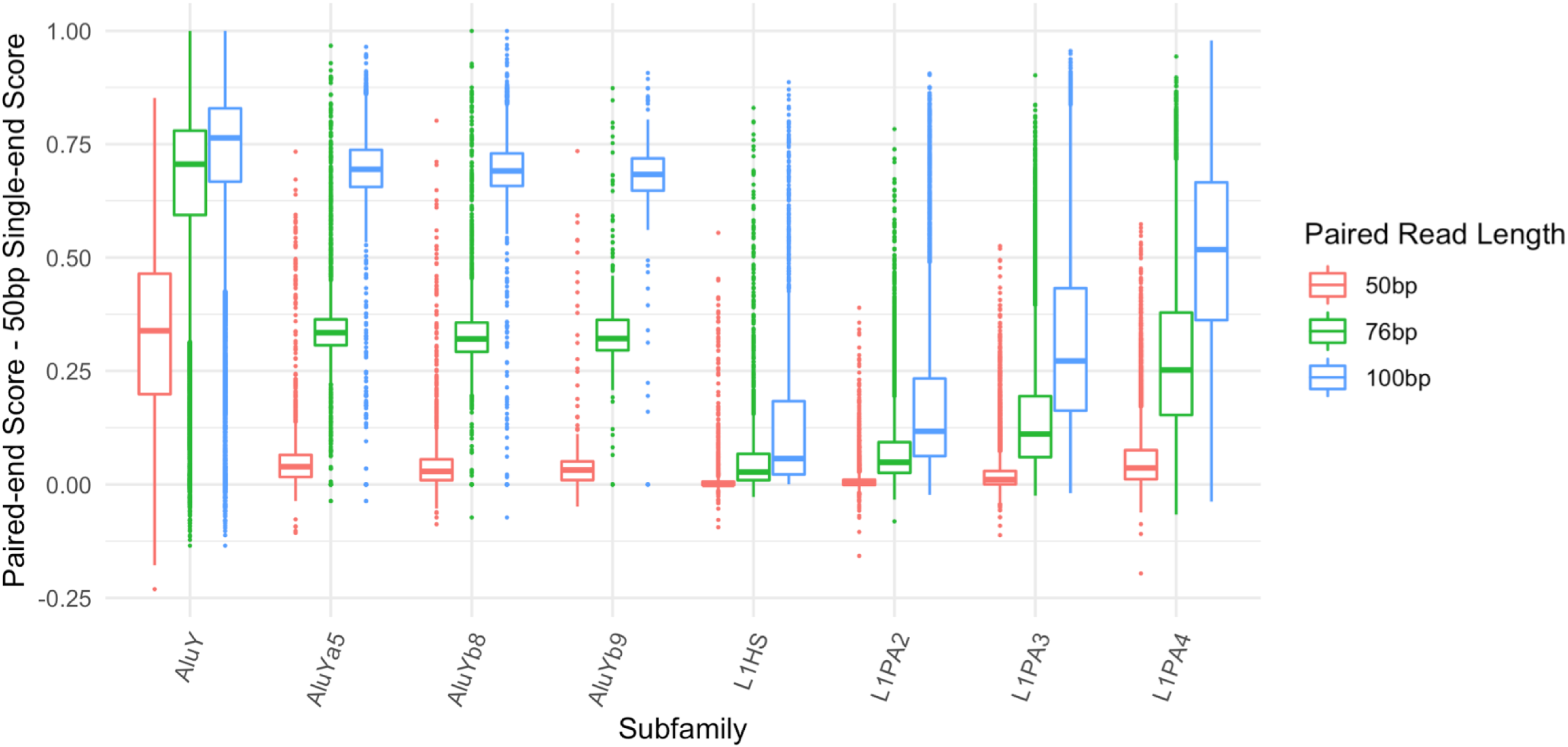
Increase in mappability between single-end and paired-end reads for low mappability families. For 8 of the lowest mappability scoring subfamilies in L1 and Alu, the boxplots show the difference between paired-end scores (at 50bp, 76bp and 100bp read lengths) and the 50bp single-end scores for every element in the subfamily.

### Effect of Read Lengths on Mappability of Elements

To investigate the effect of read length on mappability scores, we plotted the mappability scores for TE loci using our six different read libraries in Figure 6. The loci are sorted based on the 50bp single-end mappability scores. As expected, there is a strong pattern of higher mappability scores with longer read length.

**Figure 6:**
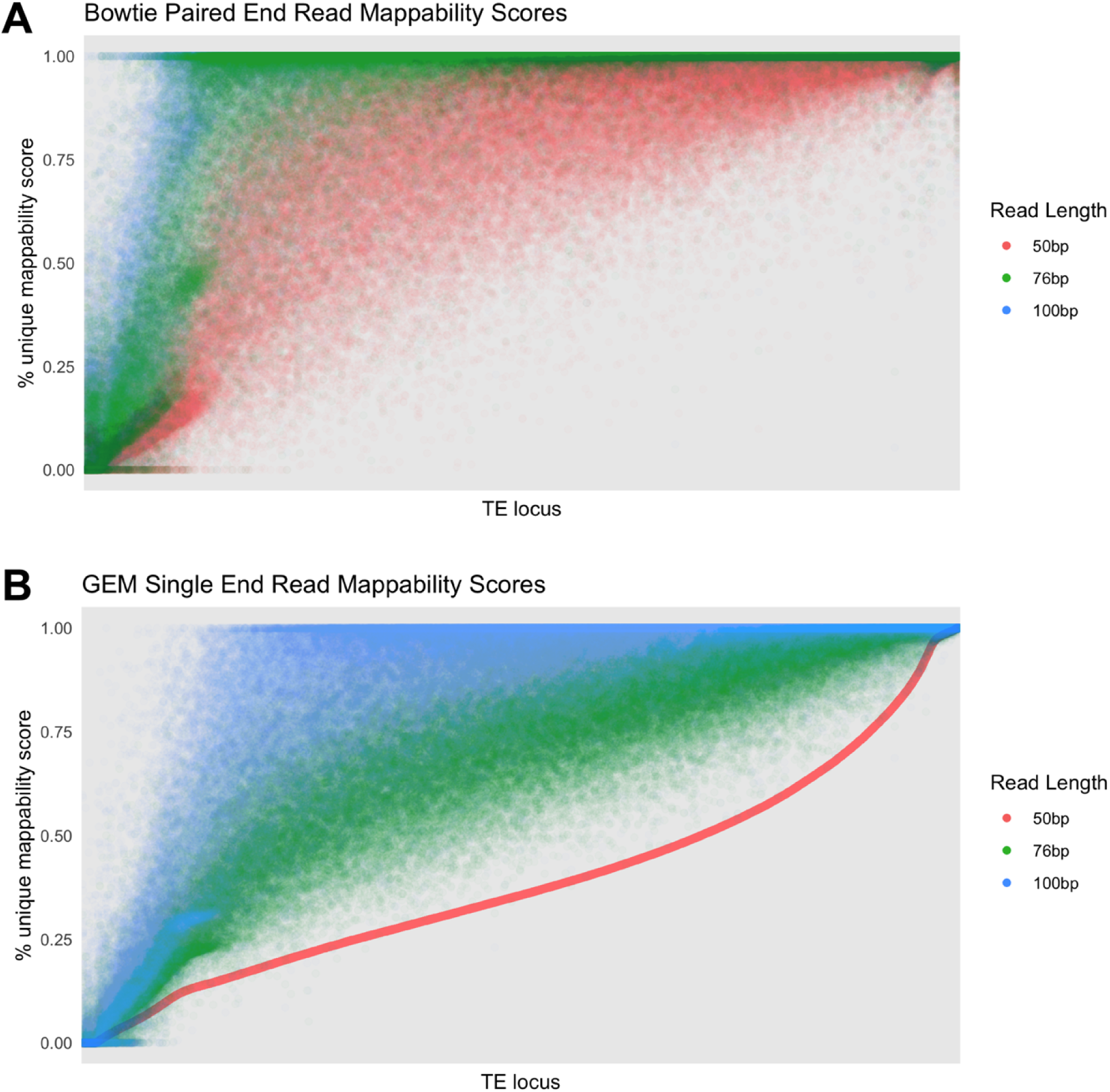
Paired-end and single-end mappability Scores for different library sizes. The mappability scores for each locus calculated using (a) paired-end and (b) single-end libraries at three different read lengths: 50bp, 76bp, and 100bp. Loci along the x-axis in both (a) and (b) are sorted in ascending order for the 50bp single-end library scores.

To further understand the gain between different library types, we plotted the differences between all paired-end and single-end library mappability scores. As shown in Figure 7, the scores from the 50bp paired-end library are very comparable to a single 76bp read library and are in general lower than a single 100bp read library. Similarly, the 76bp paired-end library has very similar scores when compared to a 100bp single-end read library. This suggests that the gain in mappability from longer single reads may outweigh the gain from a shorter paired-end read library. It is however important to note that these scores are unique to our specific definition of mappability and therefore we cannot conclusively state at what point a single-read library is more advantageous than a paired-end library for all experiments.

**Figure 7:**
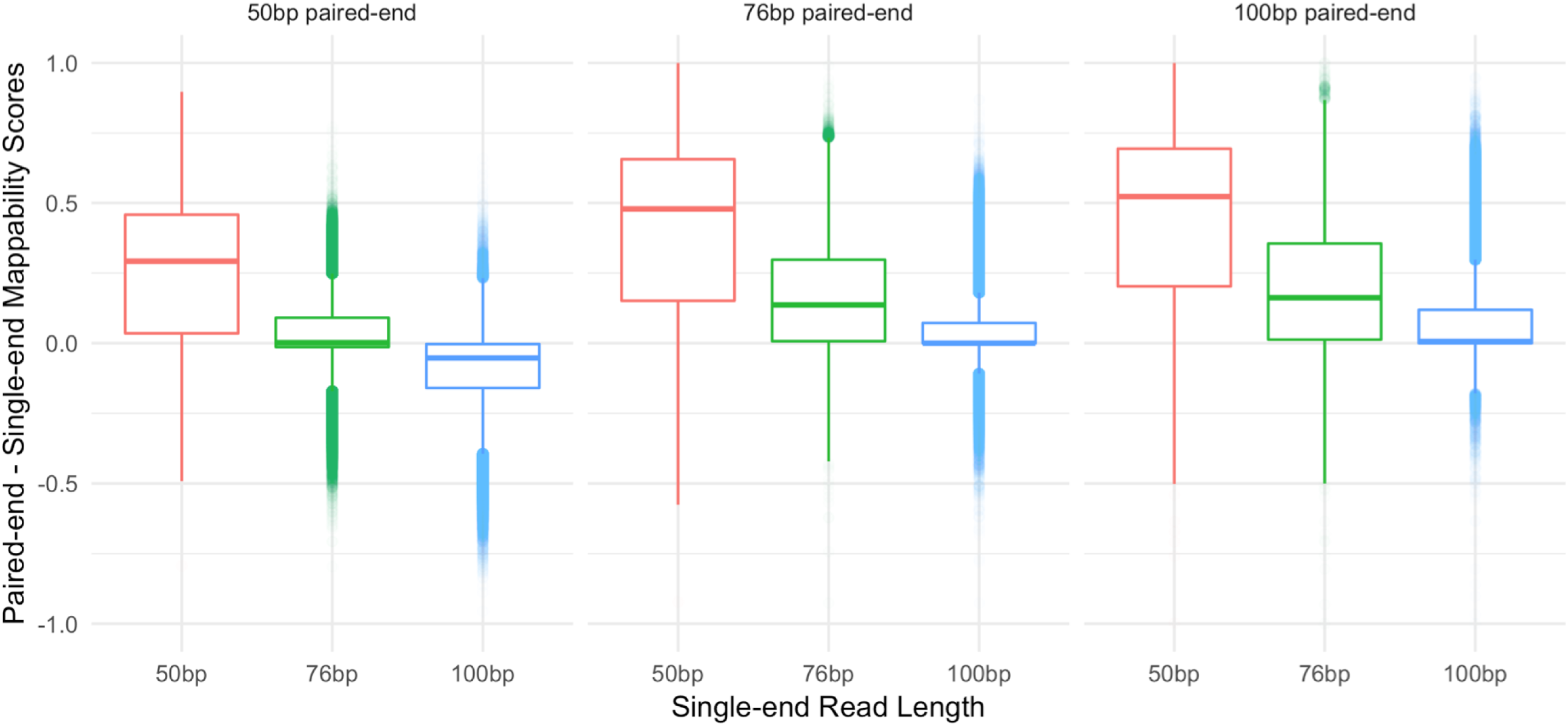
Difference in paired-end and single-end mappability scores across combinations of library sizes. Boxplots show the distribution of the differences between paired-end mappability scores (specified above the boxplots) and single-end mappability scores (specified along the x-axis) for each combination of 50bp, 76bp, and 100bp paired and single-end libraries. The scores shown here exclude TE loci where all three paired-end libraries have 100% unique scores and where length < 300bp in order to allow for a fair comparison with the 100bp paired-end library. Points < 0 signify a higher single-end than paired-end mappability score and points > 0 signify a higher paired-end than single-end mappability score.

### Mappability of L1HS

As L1HS elements are active in both germline and somatic cells (17,18), their mappability is of particular interest. Figure 8 shows that mappability scores for the 76bp paired-end library correlate generally with the time since the L1 elements were active in the human genome; younger subfamilies contain less mappable loci than older subfamilies. This is expected because over time sequences acquire variants and therefore become more unique leading to higher mappability scores. Because L1HS is the youngest subfamily of L1 in the human genome, it contains the smallest amount of uniquely mappable elements.

**Figure 8:**
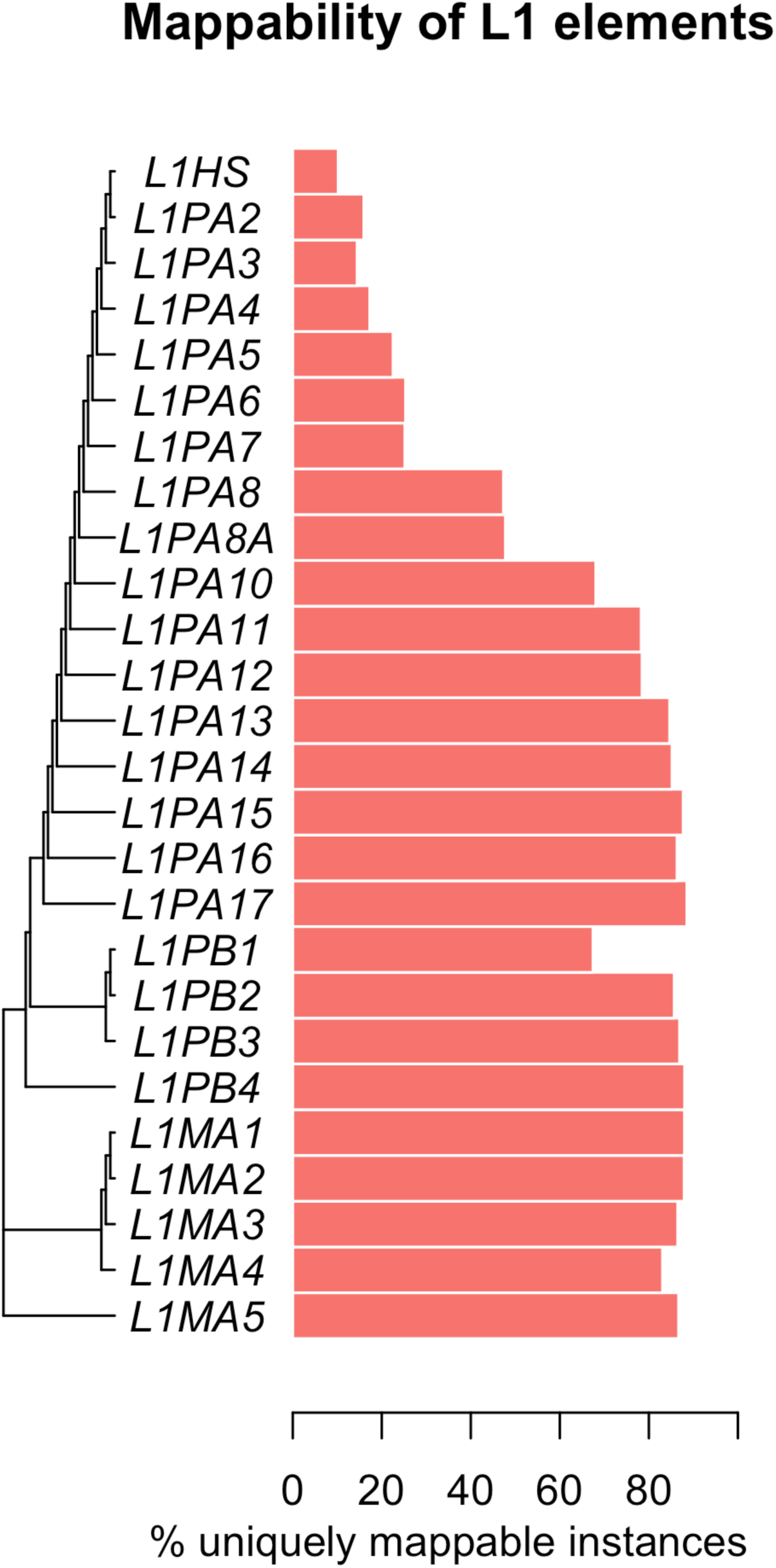
L1 phylogeny and mappability scores. For each subfamily in the phylogeny, the bar represents the percent of completely uniquely mappable loci in that subfamily according to Bowtie paired-end mappings allowing for 3 mismatches.

To further understand the mappability of L1HS, we investigated each L1HS sequence. The insertion mechanism of L1HS initiates from the 3’ end of the L1 RNA and often fails to reach the 5’ end (19) and therefore we expect more elements present and lower mappability scores at the 3’ end of L1HS consensus sequence. We used BLAST to align each L1HS locus in the genome to the consensus sequence obtained from RepBase (20) and recovered the mappability score for each locus at each position of the consensus sequence. Using those mappability scores, the trend is confirmed, shown in Figure 9. The median mappability score across loci decreases and the number of L1HS loci is greater towards the 3’ end of the consensus sequence. Conversely the median mappability increases and the number of L1HS loci is lower towards the 5’ end of the consensus sequence. For the 5’ end of L1HS, about 15% of the elements have a mean mappability greater than 0.20, meaning, the source of the read can be narrowed down to five potential L1HS loci in the genome.

**Figure 9:**
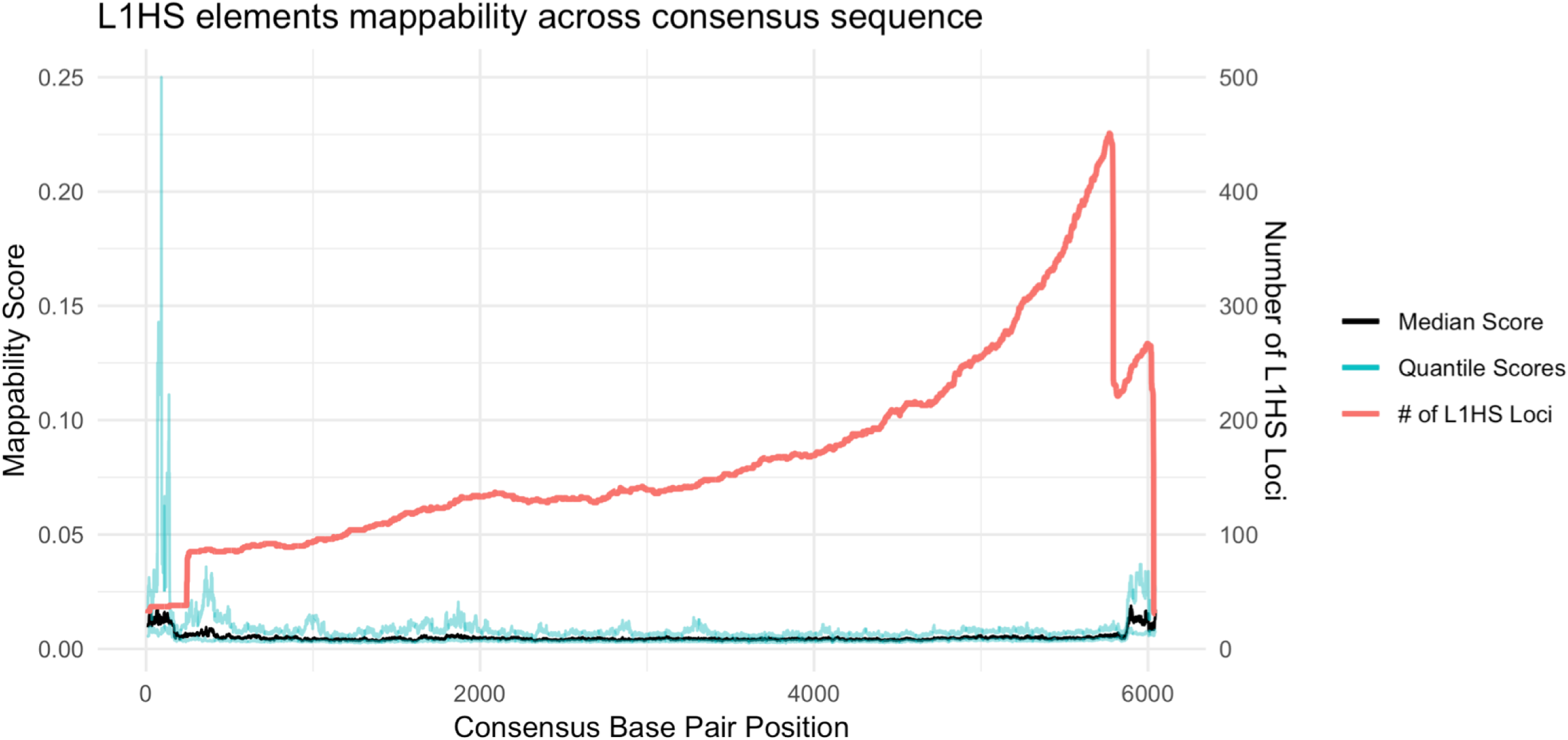
Mappability along the L1HS consensus sequence. The black line is the median mappability scores across L1HS elements and the blue lines are the 1st and 3rd quartile scores. The x axis represents the base positions along the consensus sequence. The red line is a density line of the number of loci present at that base position of the consensus sequence.

We also determined mappability scores for putatively active L1HS elements defined in L1Base 2 (21) as compared to other elements in the L1HS subfamily. These elements are classified as putatively active if they are either full-length or contain an intact ORF2 region. Figure 10a shows the distribution of mappability for both the 144 putatively active L1HS elements and the 1,542 L1HS elements not predicted to be active. The means of these distributions are significantly different (p = 2.2e-16; permutation test (22)) which corroborates the hypothesis that active elements are less mappable than inactive elements.

**Figure 10:**
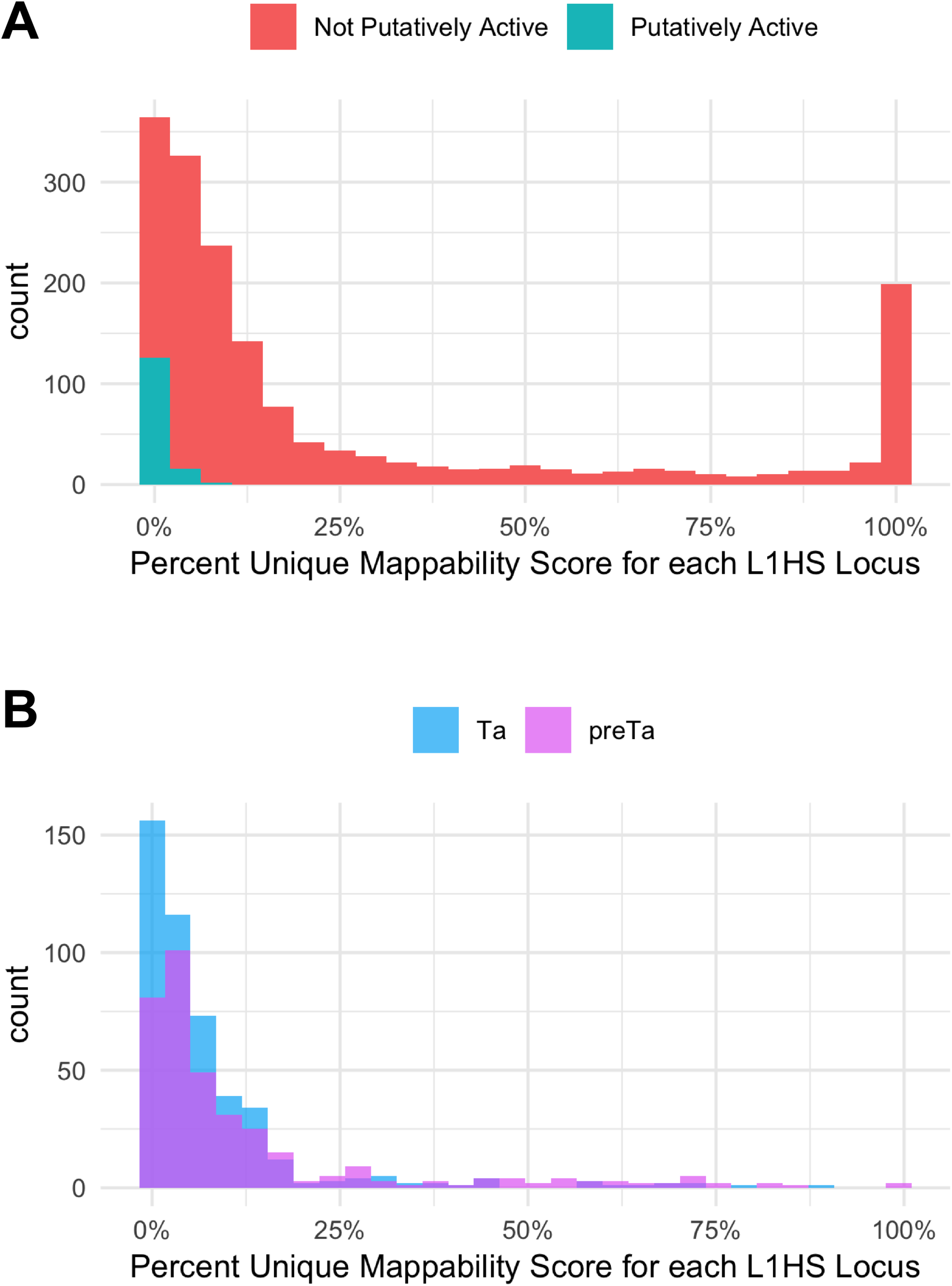
L1HS mappability score distributions. We calculated the percent of each element’s sequence with unique mappability scores and then generated a histogram from those percentages. In (a), the pink distribution is the 1,542 not putatively active elements and the blue distribution is the 144 putatively active elements. Using a permutation test, the means of these distributions are significantly different (p = 2.2e-16). In (b), the blue distribution is made of the L1HS-Ta elements and the purple distribution is made of the L1HS-preTa elements. The means of these distributions are also significantly different (p = 2.491e-06) when using a permutation test.

In addition, we further classified the L1HS elements classified in Repeatmasker into sub-clades by their characteristic 3’ UTR variants: ACA for L1HS-Ta and ACG for L1HS-preTa (23). The resulting distributions as shown in Figure 10b are also significantly different (p = 2.491e-06) again validating the idea that older elements, L1HS-preTA in this case, have higher mappability scores than younger elements, L1HS-Ta elements.

## Discussions

In our mappability analysis the insert size chosen to simulate the paired-end kmers was fixed, whereas in a true dataset the insert size would follow a distribution. However, by aligning the kmers with Bowtie’s default allowance of a variable insert size, we allow for a distribution of possible alignments. A limitation of our study is that we did not account for the polymorphic TEs present in the human population which could lower the mappability even further for younger elements. This limitation could be addressed in the future, once we have a compilation of the full-length sequences of longer polymorphic TEs utilizing long read sequencing platforms, such as the Pacific Biosciences (PacBio) platform or the Oxford Nanopore platform. Another limitation of our study is that we do not consider splicing events that happen for certain endogenous retroviruses, *e*.*g*.HERV-K(HML-2), or MLT2A1 LTRs. This is a common limitation found in virtually all mappability studies due to feasibility issues, including ones aimed at regular genes. For example, for genes, artificial transcripts are generated by concatenating all exons within the boundary of the gene, and mappability is calculated based on the artificial transcript, ignoring realistic splicing events observed. We partially address this issue by examining the alignment using STAR, a splicing oriented alignment software. But, we found that in general Bowtie gave us more conservative mappability, *i*.*e*. STAR was missing a lot of mappable positions.

In conclusion, we present our findings that increased read length and paired-end libraries result in a rise in the mappability of repeat elements, to the point where a majority of elements are uniquely mappable using Bowtie with paired-end reads allowing for 3 mismatches. Additionally we predict that a longer read library size of 150 bp would further increase mappability scores, although we have only calculated scores for up to 100bp paired-end reads for feasibility reasons.

Based on previous studies (24), only a few of all TE loci in the genome have been observed to be transcriptionally active depending on the tissue and developmental time point, and the majority of the uniquely mappable TE loci we have identified may be biologically irrelevant. Even so, these mappability estimates give us a set of references that we can utilize to identify the few transcriptionally active and potentially biologically relevant loci with confidence. Overall, our paired-end mappability analysis suggests that longer paired-end read libraries can be confidently mapped to repetitive regions and specifically to the locus-level of the majority of TEs.

## Abbreviations

TE: Transposable Element
GEM: GEnome Multitool
GTEx: Genotype-Tissue Expression Project

## Declarations

### Ethics approval and consent to participate

Not applicable.

### Consent for publication

Not applicable.

### Availability of data and material

Six hg38 mappability tracks (GEM Mappability single-end 50bp, 76bp, and 100bp allowing 3 mismatches, Bowtie paired-end 50bp, 76bp, and 100bp allowing 3 mismatches) are available for upload as a custom track on the UCSC browser at https://github.com/HanLabUNLV/TEmappability (Supplemental Figure 4), as well as locus level mappability scores for all TEs in the RepeatMasker database.

### Authors’ contributions

CES and MVH conceived the study, ran the analysis, and wrote the manuscript. Both authors critically reviewed and approved the final manuscript.

## Acknowledgements

Not applicable.

