## Supplementary Materials for "Paired-end Mappability of Transposable Elements in the Human Genome"

**Additional Files**

*Supplemental Figure 1:* Colored by class distinction, each bar represents the percent unique elements in each subfamily. An element is only considered unique if every position in its sequence has a unique mappability score. Both the Bowtie and STAR scores were generated by 76bp paired-end simulated kmers.

*Supplemental Figure 2: Bowtie mappability calculation method comparison*

Comparison of mappability scores when calculated with two different methods. Method 1: For a single base position, if a kmer mapped to that position and jellyfish (the kmer generator) reported that kmer to appear 5 times in the genome, the mappability of that position would be 1/5. Method 2: Identify a kmer which maps exactly to a base position and take the inverse of how many times that kmer appears at other positions in the bamfile. Although the figure does not reflect it very well, over 90% of the points are placed along the diagonal of the plot. In this study we use method 1, the more conservative measure, for all paired-end score calculations.

*Supplemental Figure 3: Bowtie vs GEM Mappability scores for single-end mappability*

Comparison of Bowtie 76bp mapping and GEM Mappability 76-mer mappability scores with 0, 1, and 2 mismatches.

*Supplemental Figure 4: UCSC Genome Browser tracks*

Visualization of the 6 new hg38 UCSC Genome Browser tracks. The paired-end scores are in blue and the single-end scores are in red.

*Supplemental Table 1: TE class and family mappability scores*

Percent of elements in each TE class and family with completely unique mappability scores based on Bowtie alignment of 76bp simulated reads allowing for 3 mismatches.

*Supplemental Table 2: Ancient TE families with unusually low mappability and accompanying segmental duplications.*

This table enumerates a number of ancient TE families which have unusually low mappability scores. Though some of these scores can be accounted for by recent segmental duplications, there still remains a substantial portion of older elements with low mappability scores that could be furthered explored.

Supplemental Figure 1

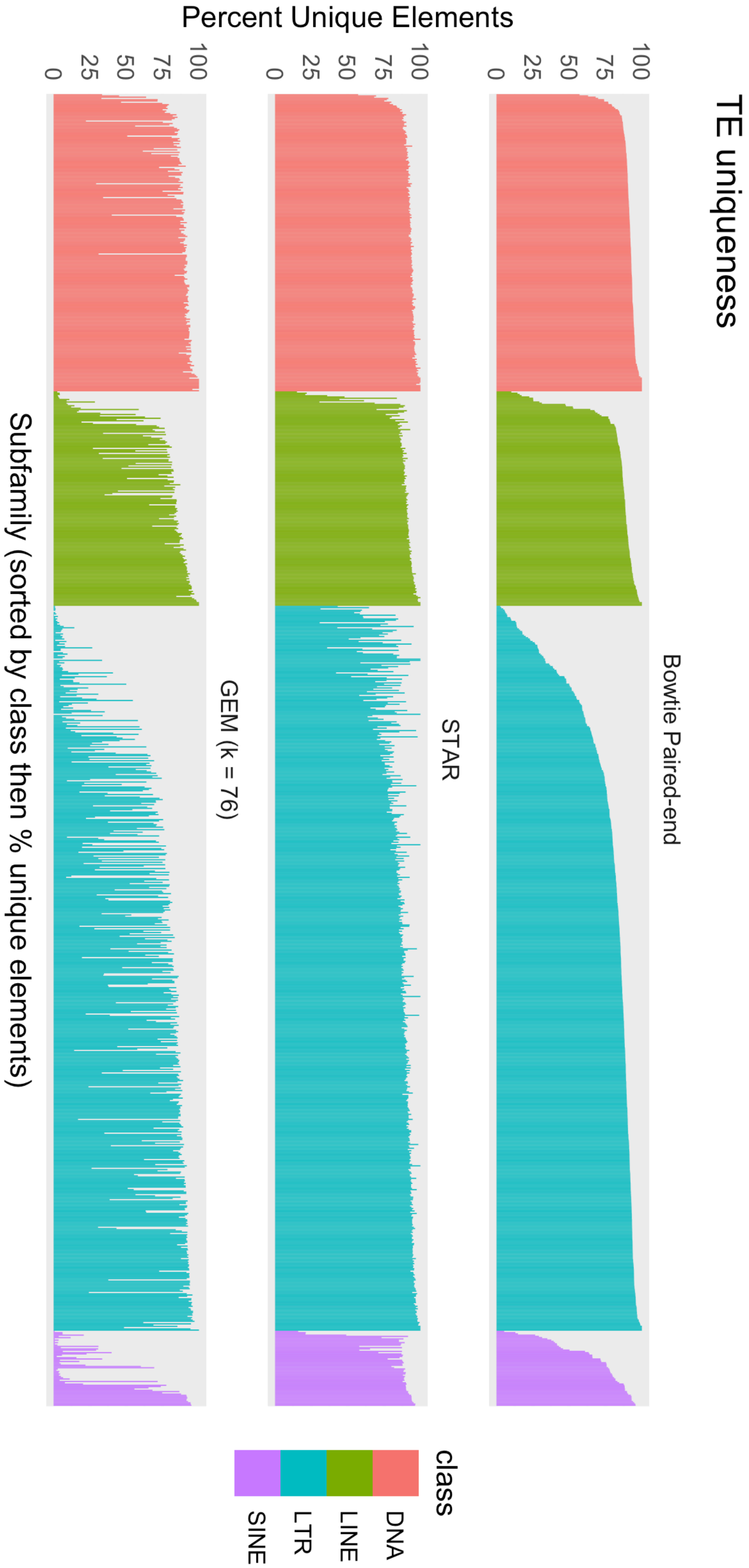

Supplemental Figure 2

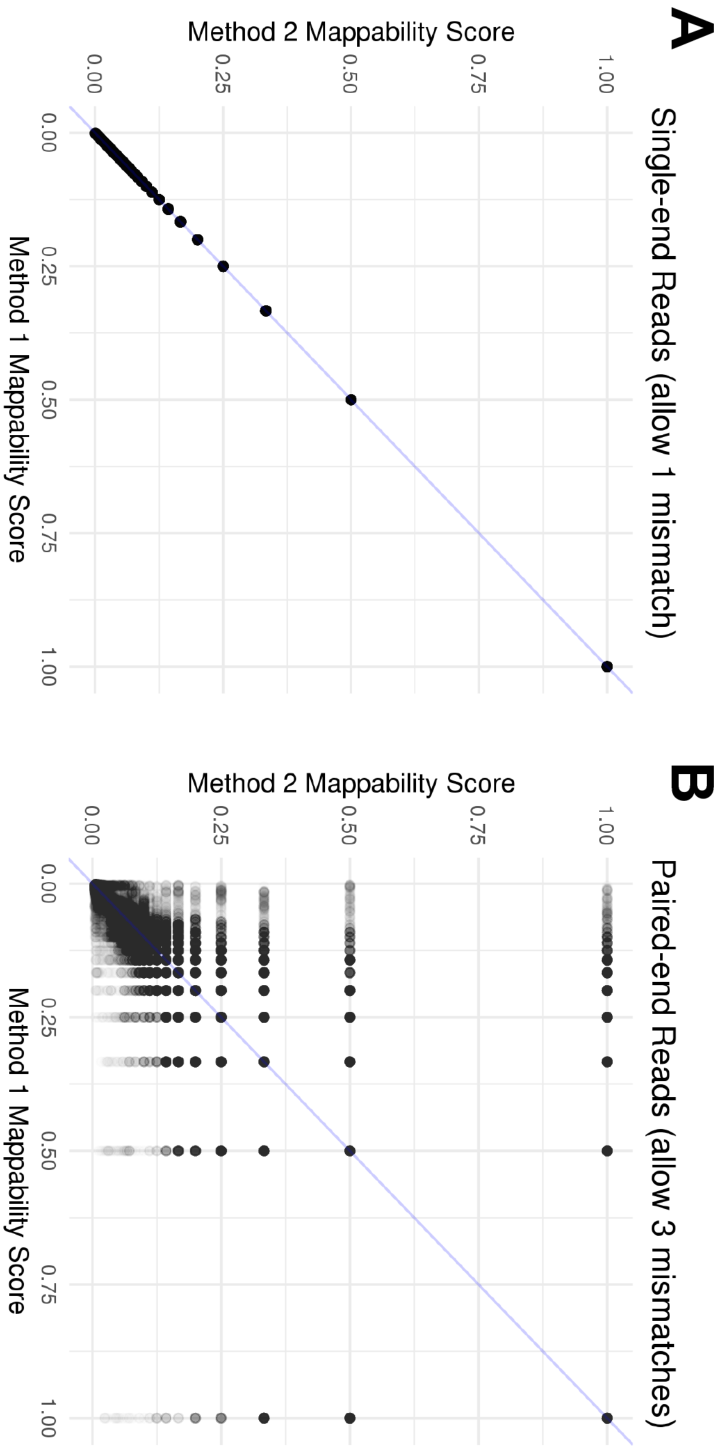

Supplemental Figure 3

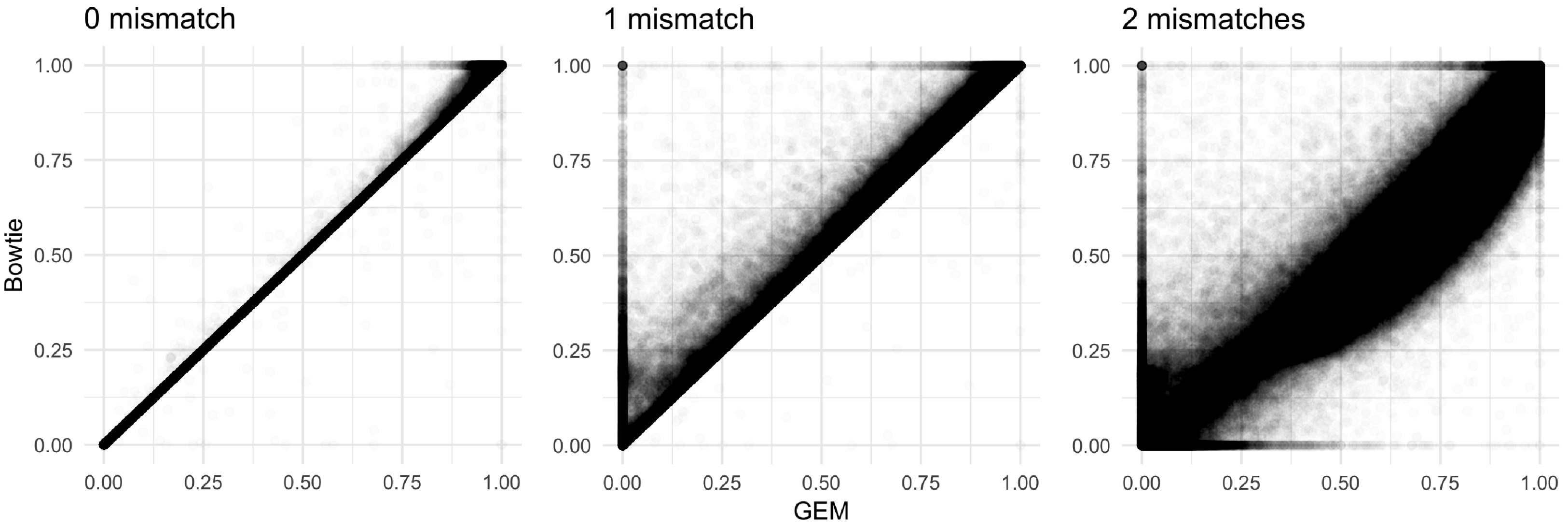

Supplemental Figure 4

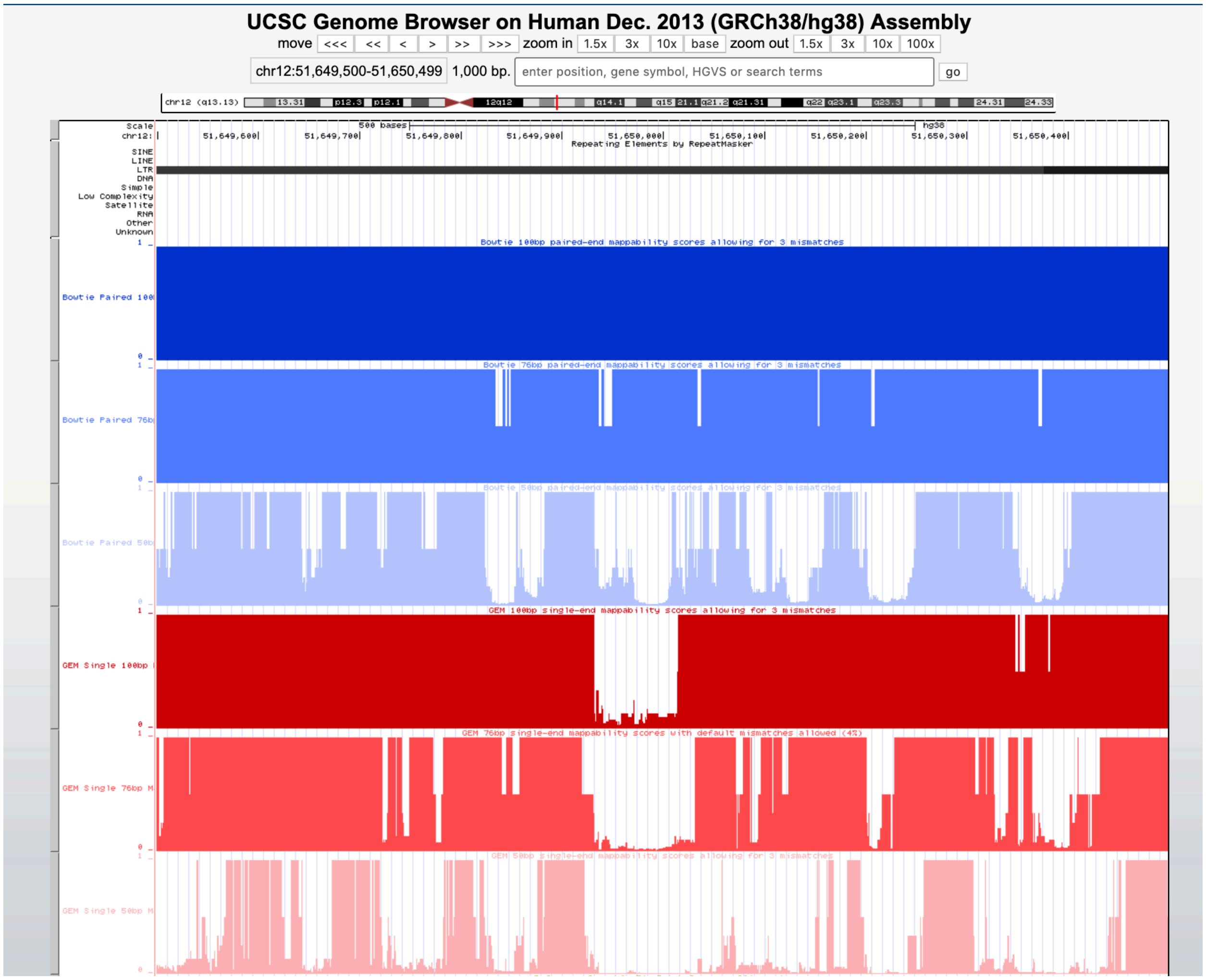
